# A Novel Fluorogenic Spin Probe for Real-time Monitoring of Mitochondrial Reactive Oxygen Species (mROS) Generation in an *in vitro* Stroke Model

**DOI:** 10.1101/2023.09.17.558106

**Authors:** Shanshan Hou, Yue Bi, Chunyan Li, Zhiying Shan, Lanrong Bi

## Abstract

The extent of oxidative damage caused by a stroke can be assessed by monitoring mitochondrial reactive oxygen species (mROS), which can further aid in early diagnosis and evaluation of treatment efficacy. Mitochondrial ROS have been identified as a potential biomarker for stroke, as they are known to cause oxidative stress and mitochondrial dysfunction in brain tissue. However, monitoring reversible mROS levels has been challenging due to the lack of appropriate probes. In this study, we developed a fluorogenic spin probe, Mito-RT-ROS, based on the rhodamine skeleton, which demonstrated good sensitivity for reversible monitoring of mROS in a physiological environment such as starvation and recovery. We also evaluated the probe’s efficacy in an *in vitro* stroke model for real-time tracking of mROS generation in a pathological setting.

## INTRODUCTION

Ischemic stroke happens when a sudden obstruction of blood flow to a part of the brain causing a shortage of oxygen and nutrients. As a result, the neurons in the infarct core, the region directly impacted by the lack of blood supply, undergo necrotic cell death, where cells die due to the severe deprivation of oxygen and nutrients. In the case of ischemic stroke, there are two main acute treatments: intravenous thrombolysis and endovascular thrombectomy. However, both treatments are time-sensitive and must be administered within 4.5-6 hours from the beginning of the symptoms. ^1,2^ The primary limitation of these treatments is that they are most effective when administered early, and many stroke patients do not reach the hospital in time to receive them. Moreover, not all strokes are caused by large blood clots that can be easily removed or dissolved.

The penumbra refers to the region surrounding the infarct core. It comprises neurons still alive but at risk of dying if blood flow is not restored promptly. While reperfusion injury is essential for saving brain tissue, it can also cause additional damage. Several mechanisms contribute to reperfusion injury, including the overproduction of free radicals.^3^ During ischemia (lack of blood flow and oxygen), cells in the affected brain tissue can undergo metabolic changes that lead to the accumulation of reactive oxygen species (ROS) and other free radicals. When blood flow is suddenly restored, these accumulated free radicals can react with oxygen to produce more damaging molecules. This can result in oxidative stress and further injury to brain cells.

The generation of free radicals during reperfusion exacerbates tissue damage and inflammation in the aftermath of an ischemic stroke. To mitigate reperfusion injury, researchers explore various strategies, including using antioxidants and other neuroprotective agents, to help minimize oxidative stress and inflammation during the reperfusion phase of treatment.^4,5^ Currently, no approved neuroprotective agent reliably prevents or mitigates this secondary damage in the penumbra following an ischemic stroke. Developing effective neuroprotective agents for ischemic stroke is a complex challenge, and numerous experimental treatments have been explored over the years. These efforts aim to protect neurons in the penumbra, reduce inflammation, and prevent further cell injury.^6^ However, finding a universally effective neuroprotective agent has proven elusive, and research in this area continues to be a critical focus in stroke medicine and neuroscience. One of the challenges in developing effective treatments for stroke is the lack of a specific and reliable biomarker for early detection. Early detection and intervention are crucial for minimizing the damage caused by stroke.

By monitoring mROS as biomarkers, it is possible to detect stroke early. Elevated levels of mROS in blood or cerebrospinal fluid may indicate ongoing brain injury, prompting timely intervention and treatment.^7-10^ Additionally, assessing mROS may help evaluate the effectiveness of stroke treatments. Measuring changes in mROS levels can be done when antioxidant medications or neuroprotective strategies reduce oxidative stress. Understanding the role of mROS in stroke pathology can facilitate research into new treatment strategies. By monitoring these radicals, early diagnosis, treatment assessment, and the development of targeted stroke therapies can be facilitated, ultimately improving patient outcomes.

Currently, only a few fluorescent probes are available to detect oxidative stress. One commonly used probe is 2’,7’-Dichlorofluorescin diacetate (DCFH-DA), which helps detect ROS within cells. However, DCFH-DA has some limitations that need to be considered. For example, DCFH-DA is a cell-permeable compound that enters cells and gets deacetylated by intracellular esterases to form DCFH. Unfortunately, the exact subcellular location of DCFH is not well controlled, and it can spread throughout different cellular compartments. This can make it challenging to measure ROS in specific organelles accurately. Additionally, DCFH can be non-enzymatically oxidized, producing a fluorescent product (DCF) without ROS. This background fluorescence can lead to false-positive signals. Moreover, the fluorescence of DCF generated after reacting with ROS can quickly degrade or photobleach over time upon exposure to light. This can affect the accuracy and reproducibility of measurements, especially during extended experimental procedures.^11,12^

MitoSOX Red is a fluorescent dye detecting superoxide radicals in living cells, especially in the mitochondria. Though MitoSOX is a valuable tool for studying oxidative stress and mitochondrial dysfunction, it has some limitations. MitoSOX detects superoxide radicals specifically but can also react with other ROS and reactive nitrogen species (RNS) under certain conditions. This can lead to a possibility of inaccurate results, making it essential to consider other assays and biological data while interpreting MitoSOX signals. MitoSOX’s ability to enter the mitochondrial membrane depends on the membrane potential. MitoSOX might accumulate less efficiently in depolarized mitochondria, leading to underestimating superoxide levels. This can make it challenging to interpret results when studying conditions that affect mitochondrial membrane potential. To avoid such issues, researchers must carefully control the imaging conditions, as MitoSOX is susceptible to photo-bleaching. Prolonged or intense excitation light exposure can lead to the formation of artifacts that could be interpreted as a signal. MitoSOX’s cell permeability and uptake efficiency can vary depending on the cell type and experimental conditions. Some cell types may uptake the dye more efficiently than others, leading to discrepancies in the results. In some cases, introducing exogenous fluorescent dyes like MitoSOX could interfere with cellular functions and signaling pathways, leading to unintended physiological effects. ^11-13^

Boronate-based fluorogenic probes react stoichiometrically with H_2_O_2_ to form fluorescent phenolic products. The signal intensity changes of the fluorescent phenolic products are used for monitoring intracellular H_2_O_2_. Fluorogenic boronates conjugated to a triphenylphosphonium group were also developed for tracking mitochondrial H_2_O_2_. Mitochondria-targeted boronate (Mito-B), in combination with LC-MS-based analysis, has been used to detect mitochondrial H_2_O_2_ formed *in vitro* and *in vivo*. One limitation of boronate probes is the unfavorable reaction kinetics with H_2_O_2_. In addition, other oxidants (ONOO-or HOCl) react with boronates several orders of magnitude faster, forming the same phenolic products. Thus, correlating the fluorescent signal changes of boronate probes to intracellular H_2_O_2_ may lead to an incorrect interpretation of the results.^11-13^

Fluorogenic probes offer a non-invasive way to monitor free radical activity in living systems, which is impossible with traditional electron spin resonance (ESR) spectroscopy and has limitations related to sensitivity, sample requirements, and chemical specificity.^14-16^ Also, ESR does not have the spatial resolution to detect free radicals at the levels of individual cells or subcellular structures. Previously, we have synthesized a series of rhodamine-based fluorogenic spin probes that can be used for imaging •OH in living cells.^15^ These probes are designed to be highly selective for •OH in aqueous solution, which makes them less susceptible to interference from other ROS. As a result, these probes enable efficient •OH imaging in biologically active samples. We have demonstrated that the unique features of these probes, such as their high specificity and selectivity, offer a promising alternative to previous methods for •OH detection. ^15^

In our present study, we target specific cellular compartments with a new fluorogenic spin probe, Mito-RT-ROS probe, to study localized oxidative stress and ROS production. This is particularly important for understanding the heterogeneous distribution of radicals in complex biological systems. The Mito-RT-ROS probe was designed to achieve high sensitivity and selectivity for mROS. Herein, we report the design, synthesis and application of the Mito-RT-ROS probe, for detecting mROS generation associated with stroke.

## MATERIALS AND METHODS

### Synthesis Procedure

All the chemicals were purchased from Sigma Aldrich. Unless otherwise stated, all reactions were run under a nitrogen atmosphere (1 bar). The purity (>97%) of the intermediates and the final products were confirmed on both TLC (Merck silica gel plates of type 60 F254, 0.25 mm layer thickness) and HPLC (Waters, C18 column 4.6mm x 150 mm). NMR spectra were recorded on Bruker Advance 500 spectrometers. FAB-MS was determined by VG-ZAB-MS high-resolution GC/MS/DS and HP ES-5989x.

### Synthesis of Mito-RT-ROS Probe

The Mito-RT-ROS probe was synthesized using sequential “click” reactions under mild conditions (**Scheme 1**). To prevent fluorophore/nitroxide cleavage, the nitroxide was introduced into the molecular probe through the “click” reaction. Copper(I) iodide was used as the copper(I) source instead of the more common Cu (II)SO_4_/sodium L-ascorbate system to avoid reducing the nitroxide to the nonparamagnetic hydroxylamine derivative by sodium L-ascorbate during the click reaction. All intermediates were characterized by NMR, and the structural characteristics of the target compound was confirmed by high-resolution mass spectrometry. Additionally, EPR spectroscopy was used to analyze the nitroxide-labeled probe and confirm the intact nitroxide label. The Mito-RT-ROS probe is stable and was stored at room temperature in sealed and light-protected bottles until use.

**Scheme 1.**
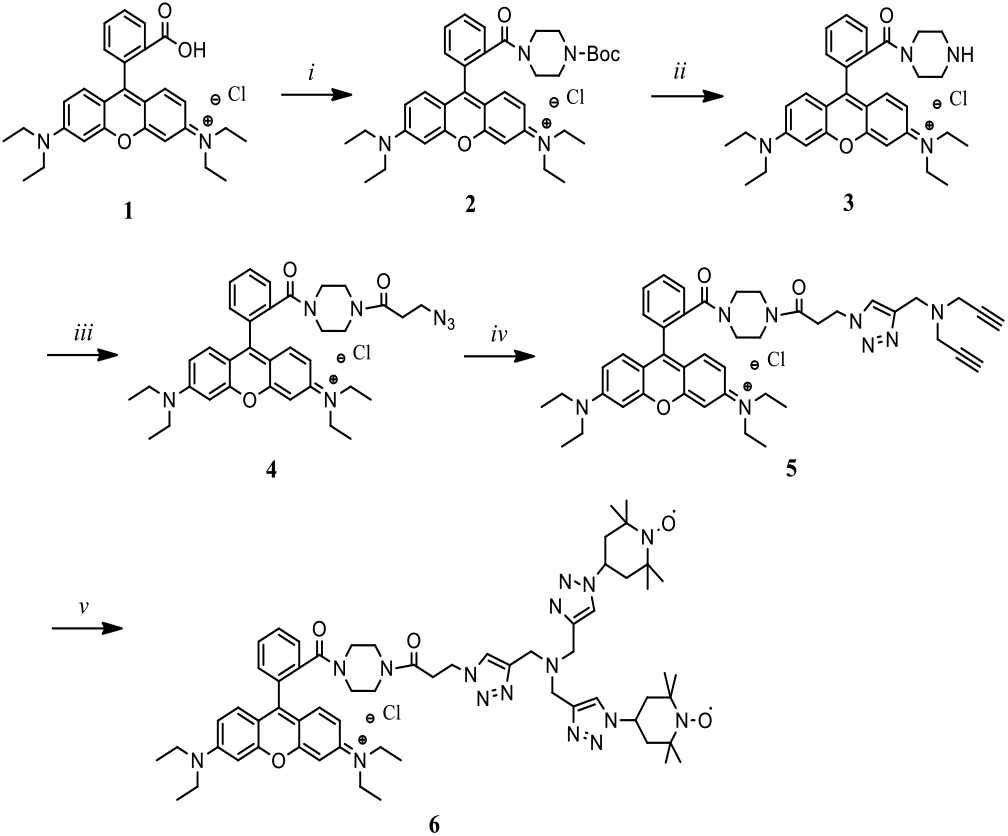
Reagents and conditions. *i*. Bocpiperazine, Et_3_N, HBTU, CH_2_Cl_2_, 2 h; *ii*. TFA:CH_2_Cl_2_1:5, 1 h; *iii*. 3-azidopropanoic acid, Et_3_N, HBTU, CH_2_Cl_2_, 2 h; *iv*. Tripropargylamine, CuI, ACN, 3 h; *v*. 4-azido TEMPO, CuI, ACN, 3 h.

#### Rhodamine B Boc-piperazine Amide 2

To a stirring solution of Rhodamine B (compound **1**, 2.00 g, 4.18 mmol), Boc-piperizine (0.78 g, 4.18 mmol) and Et_3_N (1.02 g, 10.04 mmol) in CH_2_Cl_2_ (50 mL), was added HBTU (1.90 g, 5.02 mmol). The mixture was stirred for 2 h at room temperature and diluted in CH_2_Cl_2_ (50 mL). The solution was washed with brine (50 mL×2), dried on anhydrous Na_2_SO_4_, filtered and concentrated. Flash column chromatography of the residue on silica gel (CH_2_Cl_2_:MeOH, 20:1) afforded the desired compound **2** as a purple foam (2.28 g, 91.2%).^1^H NMR (400 MHz, CDCl_3_) δ ppm 7.62-7.57 (m, 2H), 7.48-7.44 (m, 1H), 7.25 (m, 1H), 7.16-7.11 (m, 2H), 6.84 (d, *J* = 9.0 Hz, 2H), 6.69 (d, *J* = 2.4 Hz, 2H), 3.54 (h, *J* = 7.8 Hz, 8H), 3.29 (s, 4H), 3.19 (s, 4H), 1.33 (s, 9H), 1.22 (t, *J* = 7.8 Hz, 12H). ^13^C NMR (100 MHz, CDCl_3_) δ ppm 167.84, 157.87, 155.95, 155.78, 154.63, 135.23, 132.13, 130.92, 130.47, 130.26, 127.75, 114.23, 113.84, 96.44, 80.54, 46.22, 28.43, 12.70. HRMS (FAB+) *m/z* 611.3597, calcd for C_37_H_47_N_4_O_4_ 611.8046.

#### Rhodamine B Piperazine Amide 3

To a stirring solution of Rhodamine B Boc-piperazine amide (**2**, 11.01 g, 18.41 mmol) in CH_2_Cl_2_ (150 mL), was added TFA (30 mL) dropwise under ice bath. The mixture was stirred for 1 h at room temperature and adjusted to pH 8 with saturated NaHCO_3_ solution. CH_2_Cl_2_ layer was separated and water phase was extracted with CH_2_Cl_2_ (150 mL×2). The combined CH_2_Cl_2_ layers were dried on anhydrous Na_2_SO_4_, filtered and concentrated. Flash column chromatography of the residue on silica gel (CH_2_Cl_2_: MeOH, 10:1) afforded the desired compound **3** as a purple foam (6.09 g, 66.3%). ^1^H NMR (400 MHz, CDCl_3_) δ ppm 7.66-7.59 (m, 2H), 7.55-7.49 (m, 1H), 7.33-7.26 (m, 1H), 7.10 (d, *J* = 9.5 Hz, 2H), 6.86 (dd, *J* = 9.6, 2.3 Hz, 2H), 6.69 (d, *J* = 2.4 Hz, 2H), 3.71 (s, 4H), 3.63-3.57 (m, 4H), 3.52-3.48 (m, 4H), 3.10 (s, 2H), 3.02 (s, 2H), 1.26 (t, *J* = 7.1 Hz, 12H). ^13^C NMR (100 MHz, CDCl_3_) δ ppm 167.35, 157.88, 155.87, 155.82, 134.63, 131.67, 131.19, 130.60, 130.40, 130.32, 127.80, 118.56, 115.63, 114.42, 113.76, 96.51, 46.24, 44.44, 42.76, 38.57, 12.65. HRMS (FAB+) *m/z* 511.3073, calcd for C_32_H_39_N_4_O_2_ 511.6873.

#### Rhodamine B 3-azido-1-(piperazin-1-yl) propan-1-one 4

To a stirring solution of Rhodamine B piperazine amide (**3**, 2.23 g, 4.48 mmol), 3-azidopropanoic acid (0.50 g, 4.93 mmol) and Et_3_N (0.91 g, 8.96 mmol) in anhydrous CH_2_Cl_2_ (50 mL), was added HBTU (2.04 g, 5.38 mmol). The mixture was stirred for 3 h at room temperature and diluted in CH_2_Cl_2_ (100 mL). The CH_2_Cl_2_ layer was washed with brine (30 mL ×2). The organic phase was dried on anhydrous Na_2_SO_4_, filtered and concentrated. Flash column chromatography of the residue on silica gel (CH_2_Cl_2_: MeOH, 20: 1) afforded the desired compound **4** as a purple foam (0.74 g, 28.3%). ^1^H NMR (400 MHz, CDCl_3_) δ ppm 7.69-7.56 (m, 2H), 7.47 (d, *J* = 6.3 Hz, 1H), 7.32-7.22 (m, 1H), 7.15 (d, *J* = 9.5 Hz, 2H), 6.96 (d, *J* = 9.0 Hz, 1H), 6.89-6.74 (m, 1H), 6.74-6.61 (m, 2H), 3.66-3.45 (m, 10H), 3.44-3.36 (m, 4H), 3.35-3.17 (m, 4H), 2.63-2.45 (m, 2H), 1.25 (t, *J* = 7.1 Hz, 12H). ^13^C NMR (100 MHz, CDCl_3_) δ ppm 169.70, 167.98, 157.87, 155.86, 135.23, 132.20, 130.27, 127.66, 114.18, 113.94, 96.24, 94.85, 47.24, 46.23, 41.84, 41.36, 32.30, 12.72. HRMS (FAB+) *m/z* 608.3349, calcd for C_35_H_42_N_7_O_3_ 608.7636.

#### Rhodamine B 3-(4-((di(prop-2-yn-1-yl) amino) methyl)-1H-1,2,3-triazol-1-yl)-1-(piperazin-1-yl) propan-1-one 5

To a stirring solution of Rhodamine B 3-azido-1-(piperazin-1-yl) propan-1-one (compound **4**, 0.33 g, 0.57 mmol) and tripropargylamine (0.11 g, 0.85 mmol) in anhydrous acetonitrile (20 mL), was added copper iodide (17 mg, 0.09 mmol). The mixture was stirred for 3 h at room temperature, concentrated, and diluted in CH_2_Cl_2_ (50 mL). The CH_2_Cl_2_ layer was washed with brine (10 mL × 2). The organic phase was dried on anhydrous Na_2_SO_4_, filtered, and concentrated. Flash column chromatography of the residue on silica gel (CH_2_Cl_2_: MeOH, 20:1) afforded the desired compound **5** as a purple foam (0.24 g, 59.6%). ^1^H NMR (400 MHz, CDCl_3_) δ ppm 7.71 (s, 1H), 7.62 (d, *J* = 3.9 Hz, 2H), 7.48 (s, 1H), 7.34-7.24 (m, 1H), 7.20-7.12 (m, 2H), 7.03-6.77 (m, 2H), 6.68 (s, 2H), 4.59 (t, *J* = 6.4 Hz, 2H), 3.76 (s, 2H), 3.70-3.46 (m, 10H), 3.45-3.35 (m, 6H), 3.27-3.20 (m, 4H), 3.01-2.84 (m, 2H), 2.24 (s, 2H), 1.26 (t, *J* = 7.1 Hz, 12H). ^13^C NMR (100 MHz, CDCl_3_) δ ppm 168.86, 167.93, 157.88, 155.88, 135.25, 132.17, 130.25, 127.69, 124.59, 114.68, 113.95, 96.22, 79.01, 73.66, 48.02, 47.89, 46.26, 45.42, 42.15, 41.83, 41.24 33.15, 12.76. HRMS (FAB+) *m/z* 739.4084, calcd for C_44_H_51_N_8_O_3_ 739.9407.

#### Rhodamine B diTEMPO derivative 6

To a stirring solution of Rhodamine B 3-(4-((di(prop-2-yn-1-yl) amino) methyl)-1H-1,2,3-triazol-1-yl)-1-(piperazin-1-yl) pro-pan-1-one (compound **5**, 120 mg, 0.17 mmol) and 4-azido TEMPO (83 mg, 0.42 mmol) in anhydrous acetonitrile (10 mL), was added copper iodide (8 mg, 0.04 mmol). The mixture was stirred for 3 h at room temperature, concentrated, and diluted in CH_2_Cl_2_ (50 mL). The CH_2_Cl_2_ layer was washed with brine (10 mL × 3). The organic phase was dried on anhydrous Na_2_SO_4_, filtered and concentrated. Flash column chromatography of the residue on silica gel (CH_2_Cl_2_: MeOH, 20:1) afforded the desired compound **6** as a purple foam (0.08 g, 51.8%). ^1^H NMR (400 MHz, CDCl_3_) δ ppm 7.72 (br, s), 7.24 (br, s), 6.66 (br, s), 4.68 (br, s), 3.59 (br, s), 1.38 (br, s). HRMS (FAB+) *m/z* 1134.72, calcd for C_62_H_85_N_16_O_5_ 1133.69.

### Cell Culture and Live Cell Imaging

A rat neuroblastoma PC12 cell line was purchased from the American Type Culture Collection (ATCC) and cultured in RPMI-1640 medium containing 10% HS and 5% FBS at 37 °C in a humidified atmosphere of 5% CO_2_.

To induce nutrient starvation, the cells were incubated with HBSS for four hours after washing with HBSS. HBSS is commonly used for starvation studies and contains D-glucose and essential inorganic ions without amino acids and serum. After starvation, cells were allowed to recover under standard culture conditions.

For the simulated I/R, PC12 cells were washed with phosphate buffer solution (PBS) and incubated in a balanced salt solution, which contained 116 mmol/L NaCl, 5.4 mmol/L KCl, 0.8 mmol/L MgSO_4_, l mmol/L NaH_2_PO_4_, 0.9 mmol/L CaCl_2_, and 10 mg/L phenol red. The cells were then incubated in a hypoxia chamber (Themo Scientific, USA) using a compact gas oxygen controller to maintain oxygen concentration at 1% by injecting a gas mixture of 94% N_2_ and 5% CO_2_ for 2 hours. After hypoxia, the cells were transferred back to complete culture medium with oxygen. On the other hand, the sham-control cells were incubated in a regular cell culture incubator under normoxic conditions. Before simulated ischemia, cells were incubated with a complete culture medium containing either Tempol or LPTC5 (30 μM) under normoxic conditions for 12 hours, respectively. Next, the complete culture medium containing the drug was removed, and the cells were rinsed once with PBS. Then, during simulated ischemia for 2 hours, the cells were incubated with a balanced salt solution containing Tempol or LPTC5 (30 μM). Finally, the balanced salt solution was removed, and the cells were incubated with a complete culture medium (without drug) under normoxic conditions for 3 hours.

### Validate the sensitivity of the Mito-RT-ROS probe during starvation and recovery phases

We need to examine its sensitivity to gain valuable insights into the performance of the Mito-RT-ROS probe for detecting mROS generation under physiological conditions like starvation and recovery phases. This was done by subjecting a group of PC12 cells to starvation and recovery phases while maintaining a sham-control group under normal physiological conditions as a base-line reference. After applying the Mito-RT-ROS probe uniformly to all experimental groups, we used confocal fluorescence microscopy to quantify the fluorescence intensity of the Mito-RT-ROS probe over time. By comparing the intensity of the Mito-RT-ROS probe between the sham-control, starvation, and recovery groups, we could quantify and compare the probe’s sensitivity. To validate the sensitivity of the Mito-RT-ROS probe, we repeated this assay four times while keeping all other experimental conditions as consistent as possible to minimize confounding factors.

### The Mito-RT-ROS probe for real-time monitoring of mROS generation in a cellular model of stroke

Using a flow-through chamber, we established an in vitro stroke model by subjecting PC12 cells to a simulated ischemia/reperfusion injury (s-I/R) protocol. Before conducting in vivo imaging, we validated the ability of the Mito-RT-ROS probe to monitor mROS generation using this cellular stroke model. Our findings showed that subjecting the cells to 2 hours of simulated ischemia followed by 3 hours of simulated reperfusion resulted in the death of approximately 55% of the cells, as determined by the MTS assay.

In our present study, we used the Mito-RT-ROS probe to evaluate the effectiveness of two drug candidates in reducing the production of mROS during s-I/R. We selected Tempol^18-20^ and LPTC-5^17^ as drug candidates for in vitro testing due to their potential to reduce oxidative stress and inflammation, which are mechanisms associated with I/R injury. We divided the cells into four experimental groups: (i) the sham-control group, which was exposed to standard cell culture conditions; (ii) the simulated ischemia (s-I) group, which was subjected to 2 hours of oxygen and glucose deprivation; (iii) the simulated ischemia/reperfusion (s-I/R) group, which was subjected to 2 hours of ischemia followed by 3 hours of reperfusion (recovery of oxygen and glucose); (iv) the drug-treatment group, drugs (Tempol/LPTC-5) were given before simulated ischemia.

Before inducing simulated ischemia, Tempol/LPTC-5 was administered to determine their capacity to mitigate or reduce s-I/R as a preventive measure. Subsequently, Tempol/LPTC-5 was given during reperfusion to assess their therapeutic effects. In addition, we integrated the Mito-RT-ROS probe for real-time monitoring of the production of mROS in real-time during the experiment. We then measured and recorded the levels of mROS during both the prevention and reperfusion phases while administering these two drug candidates. Finally, we analyzed the data to evaluate the impact of each drug candidate on oxidative stress and mitochondrial function during both phases.

## RESULTS

### Intracellular localization of the Mito-RT-ROS probe

To demonstrate the organelle-targetable specificity of the Mito-RT-ROS probe in living cells, HeLa cells were incubated with the Mito-RT-ROS probe and MitoTracker for 30 minutes and imaged using a confocal microscope with excitation wavelengths of 568 nm. Cancer cells (such as HeLa cells) often exhibit altered metabolism and increased oxidative stress, leading to higher hydroxyl radical generation levels than normal cells. When the Mito-RT-ROS probe was incubated with HeLa cells, it entered the cells and interacted with the hydroxyl radicals in the cellular environment. The Mito-RT-ROS probe became strongly fluorescent within a short period of incubation. This change in fluorescence indicates that this probe has undergone a chemical or structural change due to its interaction with hydroxyl radicals. As shown in **Figure 1**, the Mito-RT-ROS probe co-localizes well with MitoTracker staining, demonstrating the high mitochondria specificity of the Mito-RT-ROS probe. Quantitative co-localization analyses validate a significant correlation based on Pearson’s coefficient (0.89).

**Figure 1.**
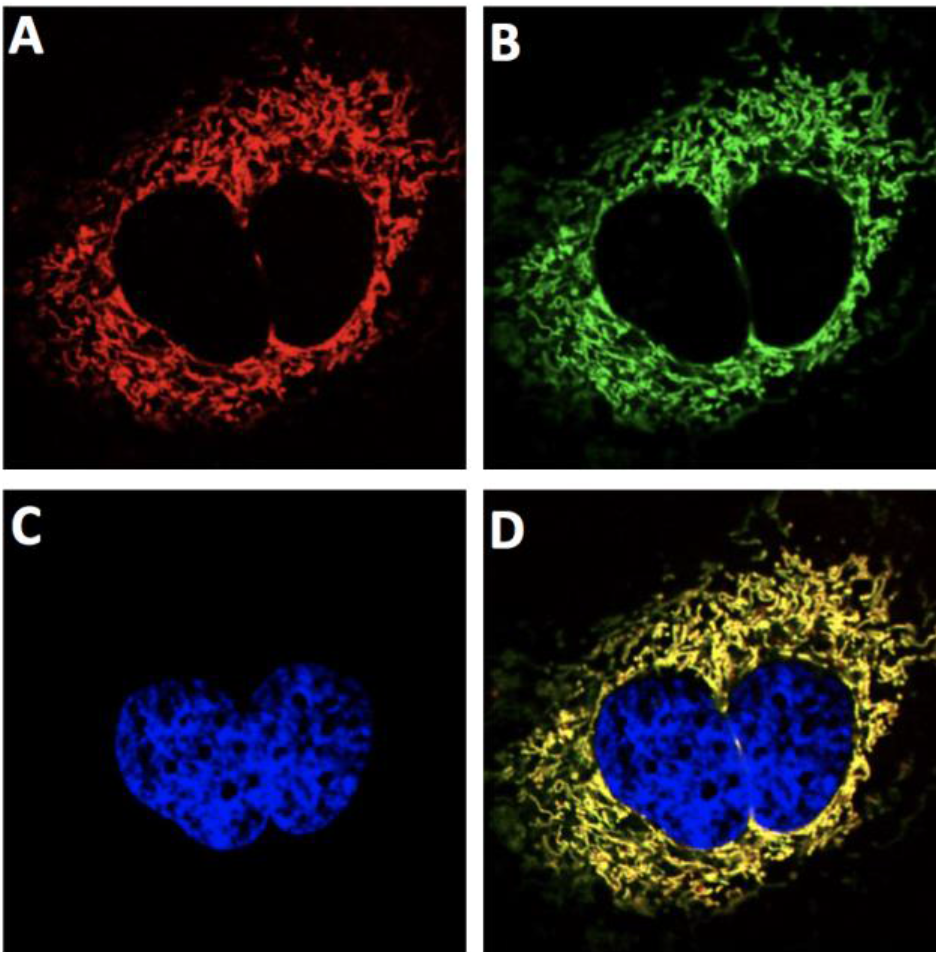
Intracellular localization of Mito-RT-ROS probe. Confocal laser-scanning fluorescent images of HeLa cells incubated with the Mito-RT-ROS probe (1μM, red fluorescence, 45 min) and then co-stained with MitoTracker (80 nM, 45 min, green fluorescence), and Hoechst 33342 (0.1 μg/mL, 30 min, blue fluorescence). The overlay images were shown in yellow fluorescence. Images were obtained with a confocal laser scanning fluorescent microscope using a 60 × objective lens in non-FBS, non-phenol red media.

**Figure 2.**
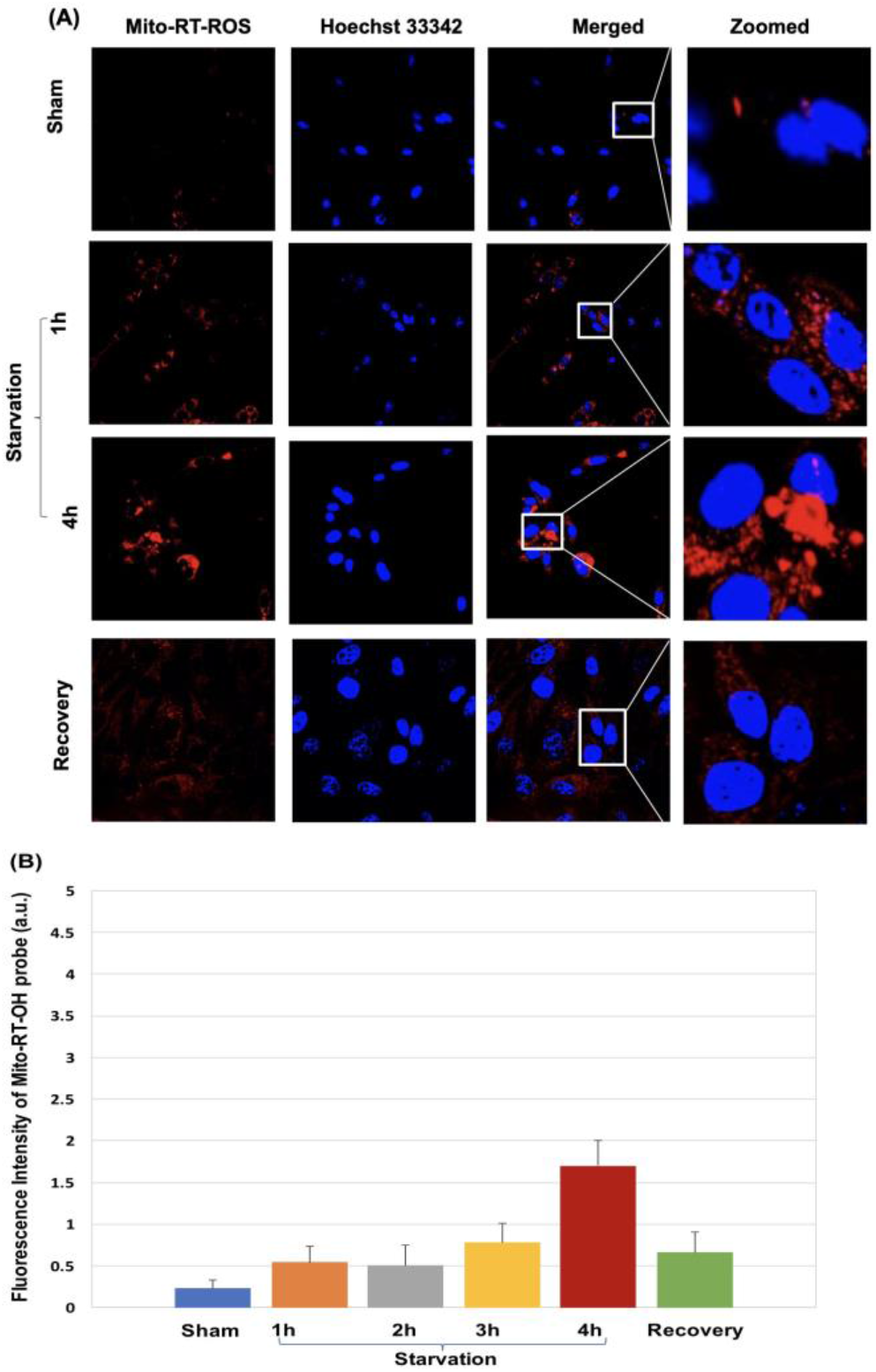
Using the Mito-RT-ROS probe for reversible monitoring of mROS generation during starvation/recovery phages. (**A**) Typical confocal fluorescence images of PC12 cells counter-stained with Mito-RT-ROS (red fluorescence), and Hoechst 33342 (blue fluorescence): The fluorescent images of PC12 cells from the sham-control group (normal cell culture condition, row 1, Fig. 2A), cells from starvation (row 2&3, Fig. 2A), and the cells from the recovery phases (row 4, Fig. 2A) are shown. (**B**) By measuring the changes of fluorescence intensity of the Mito-RT-ROS at different time points, the mROS generation was quantified throughout starvation and recovery phases. The error bars indicate the standard error of the mean from at least three independent experiments.

### The Mito-RT-ROS probe for reversible monitoring of mROS generation under a physiological setting (e.g., starvation)

Examining the sensitivity of the Mito-RT-ROS probe for mROS detection under physiological settings like starvation is crucial. Starvation is a condition where cells experience various forms of stress due to nutrient deprivation, and understanding how mROS are produced and how their levels change during such stress is essential. Monitoring mROS levels using the Mito-RT-ROS probe in starvation models may allow us to assess the degree of oxidative stress and its impact on cellular health. Elevated levels of mROS radicals are associated with various diseases, including stroke, neurodegenerative disorders, and cancer.

During a short period (1 h) of starvation, the fluorescent intensity of the Mito-RT-ROS probe increased by approximately ∼0.5 fold. This indicates that mROS is enhanced as a cellular response to nutrient deprivation stress. We observed that the fluorescence intensity of the Mito-RT-ROS probe continued to increase during extended (4h) starvation, suggesting that mROS generation is an ongoing and potentially adaptive response to prolonged nutrient scarcity. This could be related to the cell’s attempt to modulate its energy production and metabolic pathways to survive adverse conditions. When cells are returned to their normal culture conditions (i.e., nutrient-rich environment), the fluorescent intensity of the Mito-RT-ROS probe drops. This observation implies that mROS generation decreases as the cells regain access to nutrients, indicating a regulatory mechanism that responds to changes in environmental conditions.

### Using Mito-RT-ROS probe for real-time monitoring of mROS generation in an in vitro model of stroke

Before being subjected to the s-I/R protocol, PC12 cells were labeled with the Mito-RT-ROS probe. The sham-control cells displayed weak red fluorescence of the Mito-RT-ROS probe, indicating a baseline of low levels of mROS generation in the mitochondria. During simulated ischemia (s-I), most cells showed enhanced red fluorescence, limited to the mitochondria surrounding the nucleus. In contrast, simulated reperfusion (s-R) resulted in scattered and irregular staining in cells with enhanced red fluorescence. The s-I (120 min) resulted in approximately a 2.2-fold increase in mROS generation, while simulated ischemia followed by 3 hours of simulated reperfusion resulted in a 7.4-fold rise in mROS. This enhanced red fluorescence suggests that the mitochondria produce more hydroxyl radicals during the s-I/R process.

The cells pre-treated with Tempol + s-I/R had weaker fluorescence of the Mito-RT-ROS probe, suggesting a lower mROS concentration level compared to the cells from the s-I/R alone group. Surprisingly, the cells that underwent LPTC-5 + s-I/R treatment had much lower mROS levels than those from the Tempol + s-I/R or s-I/R alone groups. This implies that the elevated mROS generation caused by s-I/R could be significantly reduced with LPTC-5 intervention.

We also analyzed the changes in mitochondrial morphology in PC12 cells through co-staining with Mito-Tracker. We aimed to understand the structural changes induced by simulated ischemia and s-I/R. The changes were classified into three categories: tubular (normal), intermediate (tubular with swollen regions), and fragmented (small and globular). To determine the extent of mitochondrial morphological changes, we calculated the percentage of cells with abnormal mitochondrial morphology out of all cells with fragmented mitochondria. The quantification method involved counting 100 cells per experiment and averaging the data over four independent experiments per treatment. The sham control cells that were not treated contained mostly evenly distributed long tubular mitochondria in the cell. However, when exposed to s-I/R, the mitochondria underwent definite morphological transformations.

The appearance of mitochondria in sham-control cells was long and filamentous, as shown in **Figure 3A**. However, exposure to s-I/R caused mitochondrial fragmentation, turning tubular mitochondria into shorter, spherical ones. During the 3-hour simulated reperfusion (s-R), the number of cells with fragmented mitochondria increased significantly compared to sham-control cells. Most cells displayed fragmented mitochondria after reperfusion, which suggests mitochondrial fission and early cell death. The percentage of cells with fragmented mitochondria increased from 8.5 ± 2.8% to 79.3 ± 3.6 % with s-I/R alone. The fragmented mitochondria are mainly aggregated in the perinuclear region. Adding Tempol/LPTC-5 to the perfusate of control cells undergoing significant ischemia offered significant protection from s-I/R-induced cell death. Additionally, it significantly reduced mitochondrial fragmentation, as indicated by a significant reduction in the proportion of fragmented mitochondria observed in cells treated with LPTC-5 + s-I/R (27.3 ± 3.9 %) when compared with the cells treated with s-I/R alone (**Figure 3C**).

**Figure 3.**
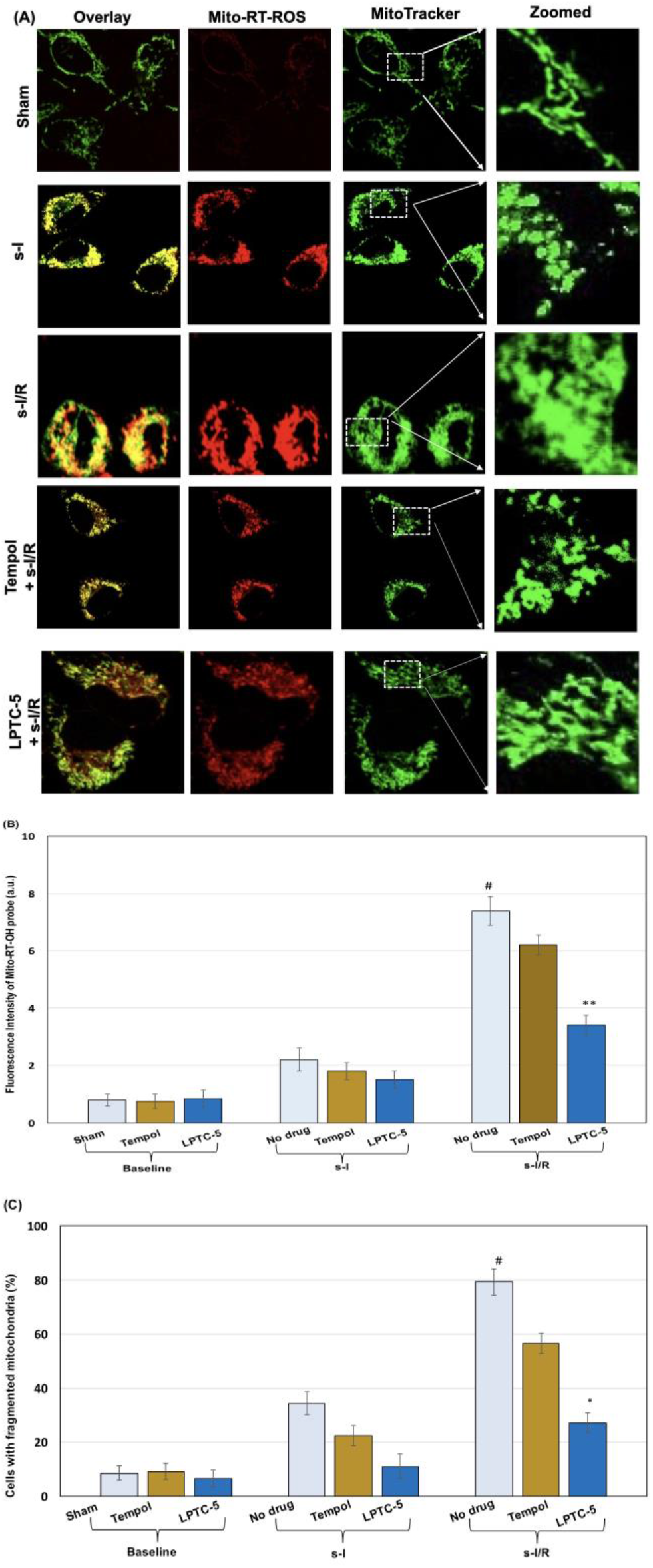
Using the Mito-RT-ROS probe for real-time monitoring mROS generation during simulated I/R (s-I/R): (**A**) Representative fluorescent images of PC12 cells counter-stained with Mito-RT-ROS (red fluorescence), MitoTracker (green fluorescence) and Hoechst 33342 (blue fluorescence): Fluorescent images of PC12 cells from the sham-control group (Sham), cells were subjected to simulated ischemia alone (s-I) or cells were subjected to simulated ischemia/reperfusion protocol alone without drug treatment (s-I/R), and with the drug pretreatment (Tempol or LPTC-5) before and during s-I/R (Tempol/LPTC-5 + s-I/R). (**B**) By measuring the changes of fluorescence intensity of the Mito-RT-ROS at different time points, the mROS generation was determined. Increases in the Mito-RT-ROS fluorescence relative to baseline values were evident during s-I/R. Pretreatment with LPTC-5 significantly attenuated the increase in Mito-RT-ROS probe fluorescence intensity during s-I/R. The fluorescence intensity of the Mi-to-RT-ROS probe was monitored to quantify the degree of the inhibition of mROS by drug treatment. The error bars indicate the standard error of the mean from at least three independent experiments (#compared with sham at baseline, *p* <0.01; **: compared with s-I/R alone, *p* < 0.01). (**C**) Cells with tubular, elongated mitochondria (%) subjected to s-I/R with/without drug treatment. (#: compared with sham at baseline, *p* < 0.05; *: compared with s-I/R alone, *p* <0.05 (s-I/R alone).

### The Mito-RT-ROS probe reveals the overproduction of mROS contributes to the release of cytochrome C from mitochondria during simulated ischemia/reperfusion

The release of cytochrome C (cyt C) during ischemia/reperfusion (I/R) injury is widely known, yet its mechanism is still unclear. We conducted a series of experiments to examine the role of mROS in cyt C release. We also investigated whether cyt C plays any part in the mitochondrial defense mechanism against oxidative stress. To this end, we first stained PC12 cells with the Mito-RT-ROS probe. We then fixed the cells at different stages, including baseline, simulated ischemia, and 3 hours of reperfusion. Finally, we immunostained the cells for cyt C to assess whether its release is related to mROS.

The sham-control cells indicate that cyt C is typically found inside mitochondria and not in the cytosol. During s-I/R, mitochondria fragmentation occurred, and cyt C was released from the intermembrane space into the cytosol. This is a response to the stress of ischemia. The release of cyt C is crucial in signaling the cell to undergo programmed cell death. It initiates a cascade of events that ultimately leads to apoptosis. The movement of cyt C throughout the cytosol plays a vital role in the cell’s programmed cell death.

We performed an immunofluorescence assay to investigate if the drug treatment could prevent the release of cyt C. **Figure 4** shows that cells that underwent LPTC-5 pretreatment retained their filamentous mitochondria. The release of cyt C from the intermembrane space into the cytosol is the opening event for the activation of mi-tochondrial apoptosis. Our results showed that LPTC-5 pretreatment significantly inhibited cyt C release during s-I/R, reducing the number of cells that release cyt C into the cytoplasm from 79.5% ± 1.8% to 21.5% ± 2.7%, as determined by cell counting (**Figure 4**). These results suggest that cyt C release during I/R depends on mROS generation. The simulated reperfusion may result in irreversible mitochondrial damage that coincides with high levels of mROS generation. Furthermore, the inhibition of cyt C release was also observed in LPTC-5-pretreated cells that underwent the simulated I/R protocol. Our findings suggest that mROS overproduction induced by s-I/R contributes to cyt C release, which could be significantly inhibited in cells subjected to LPTC-5 pretreatment.

**Figure 4.**
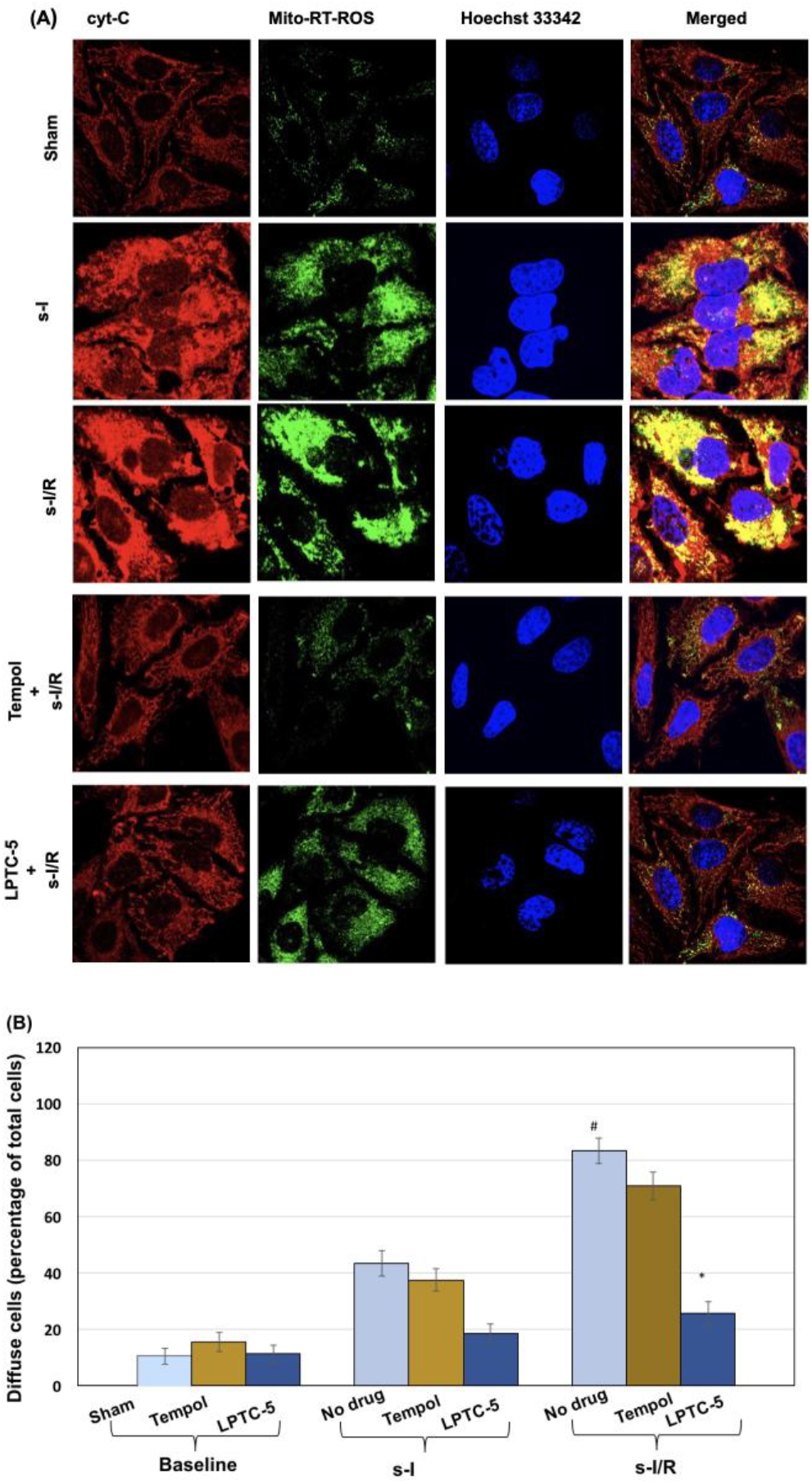
During simulated ischemia/reperfusion (s-I/R), the overproduction of mROS induces cyt C release from mitochondria. (**A**) PC12 cells were stained with the Mito-RT-ROS probe (green fluorescence). Cells were fixed at baseline, following simulated ischemia, or after 3 h of reperfusion. Cells were then immunostained for cytochrome c (red fluorescence); Representative fluorescence images of PC12 cells from the sham group (no s-I/R), cells were subjected to simulated ischemia alone (s-I) or simulated I/R injury alone without drug (s-I/R), and with drug pretreatment (Tempol/LPTC5) before and during s-I/R (Tempol/LPTC5+ s-I/R). (**B**) Cells showing redistribution of cytochrome c were quantified and expressed as a percent of the total cell population. A significant increase in the percentage of cells with cytochrome c release was observed in cells subjected to s-I/R. Cytochrome c release was significantly inhibited in cells pre-treated with LPTC. Error bars indicate the standard error of the mean from at least three independent experiments (#: compared with sham at baseline, p < 0.05; *: compared with s-I/R alone, p <0.05). The arrow indicates the redistribution of cytochrome C in the fluorescence image.

## DISCUSSION

Mitochondrial dysfunction and oxidative stress are believed to be some of the earliest events in stroke pathology. By monitoring mROS in the brain, it may be possible to gain insights into the extent and severity of oxidative damage, making them a potential biomarker for stroke. However, detecting mROS is challenging because they are highly reactive and short-lived. As a result, researchers often rely on indirect methods or assess markers of oxidative damage to infer the presence of mROS. To address this issue, researchers are exploring advanced imaging and detection techniques such as EPR spectroscopy and MRI with spin-trapping agents to indirectly detect and quantify mROS in vivo.

Currently, most fluorescent techniques typically involve the *irreversible* reaction of a non-fluorescent probe molecule with the target free radicals, producing a detectable fluorescent product. However, this “one-way” *irreversible* detection approach has limitations in responding to changes in the cellular redox environment. As a result, the chemistry of these techniques needs to be modified to provide a more dynamic response. Recently, we have designed a series of new fluorogenic spin probes that can target specific organelles, allowing us to monitor the real-time generation of the mROS. The Mito-RT-ROS probe, a “two-way” *reversible* probe, has proven effective in monitoring mROS production dynamics in a cellular stroke model.

To monitor the levels of mROS, we use fluorogenic spin probes that can be detected through fluorescence and ESR spectroscopy. ^15^ This is achieved by conjugating a nitroxide moiety with a fluorescent dye, forming a non-fluorescent compound quenched by the paramagnetic nitroxide moiety in its radical state. When the nitroxide moiety reacts with ROS, it becomes reduced and loses its paramagnetic property, resulting in fluorescence recovery of the attached fluorophore. The fluorescence signal is instantaneous and can be monitored in real-time, making it useful for capturing dynamic changes in ROS levels over short time frames, such as during cellular response to stimuli.

Examining the sensitivity of the Mito-RT-ROS probe for detecting mROS generation under physiological conditions can provide crucial information on the probe’s performance and potential applications for detecting mROS. During periods of starvation or nutrient deprivation, various biochemical processes can lead to the generation of mROS. For example, the Fenton reaction is the primary mechanism for hydroxyl radical generation. This reaction involves the interaction of iron (Fe^2+^) with hydrogen peroxide (H_2_O_2_) in the presence of oxygen (O_2_). Iron can accumulate when cells are deprived of nutrients due to altered iron metabolism. This excess iron can participate in the Fenton reaction, leading to the generation of hydroxyl radicals.

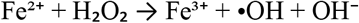

Nutrient deprivation can activate stress response pathways in cells. These pathways may involve the production of ROS, as a signaling mechanism to regulate cell survival and apoptosis. Copper ions (Cu^2+^) can also participate in reactions that generate hydroxyl radicals. Copper is a trace metal that can accumulate in cells during nutrient deprivation and catalyzes the formation of hydroxyl radicals in the presence of hydrogen peroxide.

Using the Mito-RT-ROS probe, we can continuously monitor changes in fluorescence intensity, allowing us to observe the dynamics of mROS generation. One of the notable features of this probe is its reversibility. This means that after the starvation phase, when cells enter the recovery phase, the conditions may change, and the levels of mROS may decrease. We can get an indication of this recovery process by measuring the decrease in fluorescence intensity of the Mito-RT-ROS probe.

In our present study, we employed a simulated ischemia/reperfusion (s-I/R) cellular model ^19,21^ to establish an in vitro cellular stroke model. The model mimics the lack of oxygen and nutrients to brain tissue, which is relevant to stroke pathophysiology. This model is an efficient approach to studying stroke mechanisms and potential treatments. PC12 cells exhibit neural-like characteristics and provide insights into the reactions of neuronal cells to oxygen deprivation. Using such a cellular model, researchers can investigate stroke mechanisms and screen drug candidates without the intricacy of animal models. Additionally, we can control variables like the duration and severity of hypoxia in this model to conduct standardized and reproducible experiments. By analyzing the responses of PC12 cells to hypoxia, we can gain insights into the molecular pathways contributing to cell injury or adaptation during stroke. Furthermore, cellular models of stroke can serve as a platform for testing potential therapeutic agents, speeding up the discovery of new drugs for stroke treatment. Compared to animal models, cellular models are often less expensive and require less time to set up and execute, making it possible to conduct a higher number of experiments at a faster pace.

The cells treated with Tempol/LPTC-5 and then exposed to s-I/R resulted in weak fluorescence from the Mito-RT-ROS probe, indicating that Tempol/LPTC-5 effectively reduces the generation of hydroxyl radicals in the mitochondria. This reduction in hydroxyl radical levels protected the mitochondria from oxidative damage, leading to reduced fluorescence from the Mito-RT-ROS probe. The study suggests that Tempol/LPTC-5 intervention could attenuate mROS production associated with simulated I/R. The s-I/R protocol can enhance mROS generation, leading to oxidative stress and increased fluorescence from the Mito-RT-ROS probe. Tempol, an antioxidant, can mitigate these responses to some extent by reducing mROS production. At first, it was assumed that the protection of both Tempol and LPTC-5 regarding hydroxyl radical scavenging activity should be equivalent since both compounds contain a 6-membered nitroxide moiety. However, it was found that the pretreatment of cells with LPTC-5 could significantly decrease mROS generation and provide much better protection for the cells by reducing mROS generation and cell survival. Based on this finding, we, therefore, hypothesize that the protection of LPTC-5 is not only due to its 6-membered nitroxide moiety but also due to its overall mitochondrial fusion-promoting capacity.

The release of cytochrome C is often associated with the opening of the mitochondrial permeability transition pore (MPTP).^22,23^ The MPTP is a protein complex located in the inner mitochondrial membrane. It can open under certain conditions, such as high levels of calcium ions, oxidative stress, or pH changes. Elevated levels of mROS, often seen during oxidative stress, can contribute to the opening of the MPTP. The mROS can modify and damage proteins in the mitochondria, including those regulating the MPTP. This oxidative damage can promote the pore’s opening, which may facilitate the release of cytochrome C. Once the MPTP opens, it can disrupt the integrity of the inner mitochondrial membrane, releasing cytochrome C into the cytoplasm. Cytochrome C is a critical component in the apoptotic pathway, where its release triggers a series of events that ultimately lead to cell death.

Mitochondria play a crucial role in the production of ROS, which can be harmful to the cell if not appropriately regulated. When mitochondria become dysfunctional or fragmented, they are more likely to produce excessive mROS. However, promoting mitochondrial fusion can help distribute damaged components more evenly among mitochondria, reducing the production of mROS. Moreover, mitochondrial fragmentation or damage can trigger apoptotic pathways, leading to programmed cell death (apoptosis). By promoting mitochondrial fusion, cells can maintain a more robust and functional mitochondrial network, which can help prevent or delay apoptosis, leading to improved cell survival. In addition to preventing apoptosis, promoting mitochondrial fusion can reduce the likelihood of cytochrome C release during apoptosis. Fused mitochondria create larger and more interconnected mitochondrial networks, which can dilute the concentration of cytochrome C within individual mitochondria, making it less likely to be released. Moreover, fused mitochondria are often more efficient at energy production and less stressed than fragmented ones. Healthy mitochondria are less prone to apoptotic events like cytochrome c release. Mitochondrial fusion can lead to the formation of tubular or elongated mitochondria with a more stable outer membrane. A stable outer membrane is less likely to undergo permeabilization, necessary for cytochrome C release.

## CONCLUSION

The study showed that the Mito-RT-ROS is highly sensitive in a physiological setting, particularly in cases of starvation and recovery. The Mito-RT-ROS probe was also employed in a stroke model based on PC-12 cells, demonstrating its potential use in pathophysiological settings. Compared to ESR, which is typically limited by sample constraints and requires short-term measurements, the fluorogenic spin probe reported in this study can provide continuous measurements over extended periods, making it ideal for long-term studies. Our research team is currently focused on expanding the library of these fluorogenic spin probes and will share our findings in future reports.

## LIMITATION

It is crucial to exercise caution when using mitochondrial fusion-promoting agents, as excessive fusion or dysfunction in the fusion process can adversely affect cellular physiology. Maintaining a balance in mitochondrial dynamics and function is paramount to prevent diseases associated with mitochondrial dysfunction. A comprehensive assessment is necessary to evaluate the toxicity of mitochondrial-fusion-promoting agents such as LPTC-5 fully.

On the other hand, fluorogenic spin probes are believed to be a crucial tool in distinguishing between different types of radical species and evaluating their role in oxidative stress. This is expected to have widespread applications in various domains, such as cell signaling, disease modeling, drug screening, and environmental studies. The versatility of these probes allows them to be used in experimental systems ranging from individual cells to entire organisms. Additionally, the fluorescence output of these probes can be conveniently incorporated into high-throughput screening assays, enabling the fast assessment of potential antioxidant or therapeutic compounds that regulate free radical levels. However, it is unfortunate that further testing of this probe has not been done yet.

## AUTHOR INFORMATION

## Author Contributions

^‡^Shanshan Hou and Yue Bi, these authors contributed equally.

